# FAXC Proteins of Vertebrates and Invertebrates: Relationship to Metaxin Proteins

**DOI:** 10.1101/2023.01.04.522723

**Authors:** Kenneth W. Adolph

## Abstract

FAXC proteins of vertebrates and invertebrates are shown to possess structural features similar to the metaxins, and can therefore be categorized as metaxin-like proteins. A variety of vertebrate and invertebrate species were found to encode FAXC (Failed Axon Connections) proteins. These are predicted proteins. Among the vertebrate species are human, mouse, zebrafish, and *Xenopus*. For invertebrates, FAXC proteins were most common in phyla that include Mollusca, Arthropoda, Cnidaria, and Placozoa. The presence of characteristic GST_N_Metaxin and GST_C_Metaxin protein domains was an important criterion in identifying the FAXCs as metaxin-like. In addition, the Tom37 domain was frequently found, especially in vertebrates. Also important was the possession of segments of α-helical secondary structure in a pattern of eight helices like that of metaxin 1 and other metaxins. Both the domain structures and α-helical structures are characteristic metaxin-like features. But alignment of protein sequences of both vertebrate and invertebrate FAXCs with metaxin proteins demonstrated that FAXCs and metaxins have low percentages of identical amino acids, about 20%. Therefore, FAXCs and metaxins are related, but are not the same protein. As revealed by phylogenetic analysis, vertebrate FAXCs and vertebrate metaxins 1, 2, and 3 constitute separate but related groups of proteins. Distinct phylogenetic groups were also observed for invertebrate FAXCs and invertebrate metaxins 1 and 2. The FAXC genes have neighboring genes that differ from the neighboring genes of metaxins. For vertebrates, the genes adjacent to FAXC genes are highly conserved. For invertebrates, the genes that neighbor FAXC genes are not the same as for vertebrates, but are conserved among different invertebrates. Invertebrates can have multiple FAXC genes, up to at least 12 or 13 genes. In some cases, the multiple FAXC genes exist as a cluster of genes in close proximity.

## 1. INTRODUCTION

The failed axon connections (*fax*) gene was first identified as a *Drosophila* gene that is a dominant genetic enhancer of the mutant phenotype of the *Abl* gene (Hill et al., 1995). The Fax protein is primarily expressed in central nervous system axons, as is the Abl protein (Abelson tyrosine kinase). The Fax protein is not essential for viability in otherwise wild type flies, except when flies are also mutant in the *Abl* gene. Many vertebrate and invertebrate species possess genes homologous to the *fax* gene, as investigated in this report. In humans, a *FAXC* gene is found at cytogenetic location 6q16.2. The human gene is also known as the *C6Orf168* (Chromosome 6 Open Reading Frame 168) gene. Numerous other vertebrate species have FAXC genes. Among invertebrates, phyla that include Mollusca, Cnidaria, Arthropoda, and Placozoa possess species with FAXC genes. In addition, homologous genes have been detected in fungi and bacteria (K.W. Adolph, unpublished). It can be postulated that the human FAXC protein may function in axonal development, by similarity to the *Drosophila* protein. However, the possible functions of human FAXC protein, or FAXC proteins in different species, have not been the subject of investigations reported in the literature. As a consequence, almost nothing is known about the roles of FAXC proteins in vertebrates, invertebrates, fungi, or bacteria, except for *Drosophila*.

As demonstrated in this paper, the predicted human FAXC protein and homologous predicted proteins in other vertebrates and invertebrates have structural features which characterize metaxin-like proteins. Examining these metaxin-like features is the focus of this report. Of primary importance, vertebrate and invertebrate FAXC proteins possess GST_N_Metaxin and GST_C_Metaxin domains. In addition, FAXCs and metaxins are α-helical proteins, with α-helical segments in a pattern similar to human metaxin 1 proteins and other vertebrate and invertebrate metaxin proteins. Phylogenetic analysis reveals that FAXC proteins are related to the metaxins, although FAXCs and the metaxins form separate phylogenetic groups.

Metaxin 1 and metaxin 2 of human and mouse were initially identified as proteins of the outer mitochondrial membrane with a role, experimentally determined, in the import of proteins into mitochondria. A third metaxin – metaxin 3 – has subsequently been identified, and has a wide occurrence among vertebrates including the zebrafish *Danio rerio* and the frog *Xenopus laevis* (Adolph, 2019), both commonly used research animals. For invertebrates, genes have been detected that encode proteins homologous to vertebrate metaxins 1 and 2 (Adolph, 2020a), although metaxin 3 genes have not been found. The numerous examples of invertebrates with predicted metaxin 1 and metaxin 2 proteins include the model biological research organisms *Aplysia californica* (Mollusca), *Drosophila melanogaster* (Arthropoda), and *C. elegans* (Nematoda).

Metaxin-like proteins aren’t confined to vertebrates and invertebrates. Plants also possess metaxin-like proteins. These include the model plants *Arabidopsis thaliana* and *Zea mays* (maize or corn). In addition, metaxin-like proteins are encoded in the genomes of bacteria such as *Pseudomonas aeruginosa* and *Francisella tularensis*, both causes of human disease (Adolph, 2020b). Furthermore, metaxin-like proteins are present in protists – eukaryotes that are neither animals, plants, nor fungi – and in fungi (Adolph, 2021, 2022). Examples of protists with genes for metaxin-like proteins are the model alga *Chlamydomonas reinhardtii*, the protozoan *Amoeba proteus*, and the parasite *Trypanosoma vivax*. Among the fungi are the research organism *Aspergillus nidulans* and the human pathogen *Pneumocystis jirovecii*.

Although experimental evidence has implicated a role for vertebrate metaxins in protein import into mitochondria, much more information is available for the yeast *Saccharomyces cerevisiae* (Pfanner et al., 2019; Wiedemann and Pfanner, 2017). In yeast, the SAM and TOM protein complexes work together to bring about the uptake into mitochondria of mitochondrial membrane proteins and their incorporation into the outer mitochondrial membrane. The SAM complex consists of several proteins that include Sam37 (previously Tom37), which has homology with the metaxins. The Sam37 protein possesses GST_N_Metaxin and GST_C_Metaxin domains, like the metaxins, though the major domains are the Tom37 and Tom37_C domains. The Tom37 domain is also found in many FAXC proteins.

The vertebrates and invertebrates selected for this investigation of FAXC proteins were primarily species with fully sequenced genomes. For invertebrates as well as vertebrates, obtaining genome sequences is invaluable in providing information about the proteins encoded by the genomes and the evolution of species. To be selected for this study, model organisms used in biological research also received a high priority. In addition, organisms were chosen based on their significance for human health or their economic or environmental significance. The primary vertebrate FAXC proteins in this study, in addition to human FAXC, were those of the laboratory mouse *Mus musculus* (genome sequence: Mouse Genome Sequencing Consortium, 2002), the zebrafish *Danio rerio* (sequence: Howe et al., 2013), and the frog *Xenopus laevis* (Session et al., 2016).

Among the invertebrates with FAXC genes in this study is the mollusc *Aplysia californica*, the California sea hare, which is a valuable laboratory animal for neurobiology research. Another invertebrate is *Exaiptasia diaphana*, a sea anemone of the phylum Cnidaria that is a model organism for research involving corals (genome sequence: Baumgarten et al., 2015). Cnidarians like *Exaiptasia* are the simplest animals with cells organized into tissues. The fruit fly *Drosophila melanogaster* (Phylum: Arthropoda; Class: Insecta) is perhaps the most important invertebrate model organism for genetics and developmental biology research (genome sequence: Adams et al., 2000). An additional arthropod in this study is the Atlantic horseshoe crab, *Limulus polyphemus* (Nossa et al., 2014), which has been used in medical testing for bacterial contamination. *Trichoplax adhaerens* (Srivastava et al., 2008), the simplest multicellular animal yet described, is an invertebrate of the phylum Placozoa, and is a potential model organism for cell and developmental biology research. The California two-spot octopus, *Octopus bimaculoides*, is a mollusc with a sequenced genome (Albertin et al., 2015). The nematode or roundworm *C*. *elegans* has been widely utilized in molecular and developmental biology research, particularly to study neuronal development. Its genome sequence was the first to be determined for a multicellular organism (*C. elegans* Sequencing Consortium, 1998). The starlet sea anemone *Nematostella vectensis* (Putnam et al., 2007) is a cnidarian that is a model organism in biology research. The arthropod *Hyalella azeca* (Poynton et al., 2018) is an amphipod crustacean with uses in aquatic toxicology testing.

This study was undertaken because almost nothing is known about FAXC genes and proteins. Subsequent to the Fax protein being identified in *Drosophila* as a novel protein of nerve axon bundles (Hill et al., 1995), there have been no new publications dealing with FAXC genes or the roles of FAXC proteins in any organism. However, the automated annotation of genome sequences deposited in NCBI databases has revealed that FAXC genes are present in a wide variety of organisms, both simple and complex. In addition, the annotation of the predicted FAXC proteins has suggested that the proteins have metaxin-like features, in particular conserved metaxin protein domains. The purpose of the present study was to examine more closely the existence of FAXC genes in vertebrates and invertebrates, and to investigate the relationships between FAXC proteins and metaxin proteins. The results demonstrate that FAXC genes are highly conserved among vertebrate and invertebrate species, and that the predicted FAXC proteins show a high degree of homology to the metaxin proteins. This investigation provides the basis for further research into the roles of FAXC proteins in vertebrates and invertebrates.

## 2. METHODS

The conserved protein domains of FAXCs and metaxins were identified with the CD search tool of the NCBI (Lu et al., 2020; www.ncbi.nlm.nih.gov/Structure/cdd/wrpsb.cgi). Figure 1 includes conserved protein domains provided by the CD search tool. The secondary structures (α-helical and β-strand segments) of FAXC and metaxin proteins were determined with the PSIPRED secondary structure prediction server (Jones, 1999; bioinf.cs.ucl.ac.uk/psipred/). In Figure 2, the α-helical segments of a selection of FAXC and metaxin proteins are compared by aligning the sequences using the NCBI COBALT multiple-sequence alignment tool (Papadopoulos and Agarwala, 2007; www.ncbi.nlm.nih.gov/tools/cobalt). The BLAST Global Alignment tool and the BLAST Align Two Sequences tool (Needleman and Wunsch, 1970; Altschul et al., 1990; https://blast.ncbi.nlm.nih.gov/Blast.cgi) allowed the identities and similarities of pairs of protein sequences to be detected (Figures 3 and 7). Global Align includes the full lengths of two sequences, while Align Two Sequences only includes the highest homology regions. In addition, pairs of protein sequences were aligned using EMBOSS (https://www.ebi.ac.uk/Tools/psa/emboss_needle/). Phylogenetic relationships of the proteins (Figure 4) were determined from multiple-sequence alignments using the COBALT multiple-sequence alignment tool and the phylogenetic trees produced from the alignments. Phylogenetic trees were also generated with Clustal OMEGA (https://www.ebi.ac.uk/Tools/msa/clustalo/). The genes that are adjacent to FAXC genes (Figures 5 and 6) were found using “Genomic regions, transcripts, and products” of the NCBI “Gene” database (www.ncbi.nlm.ng.gov/gene). For species with multiple FAXC genes, the same database was used to identify the multiple FAXC genes that are in close proximity (Figure 8).

**Figure 1.**
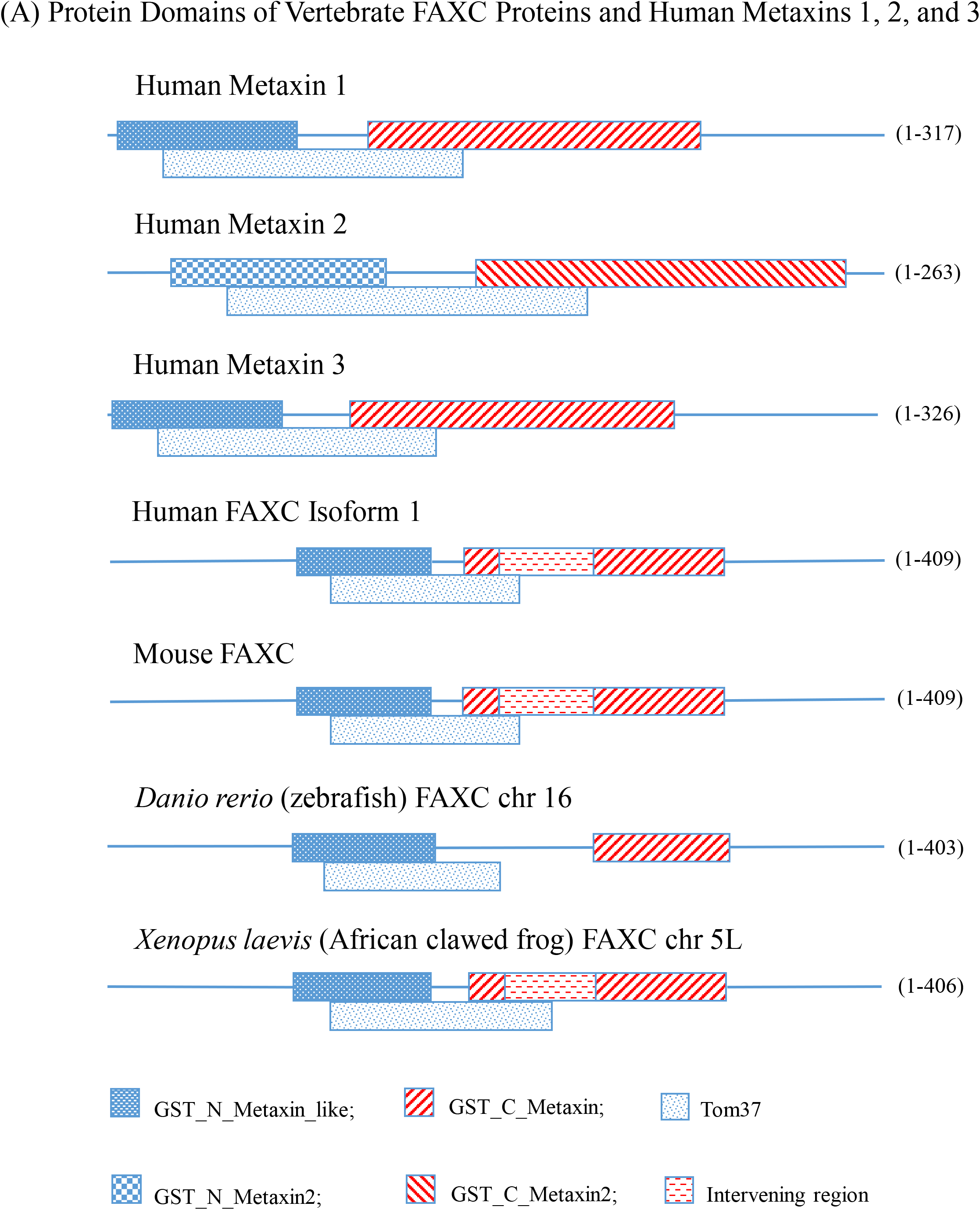
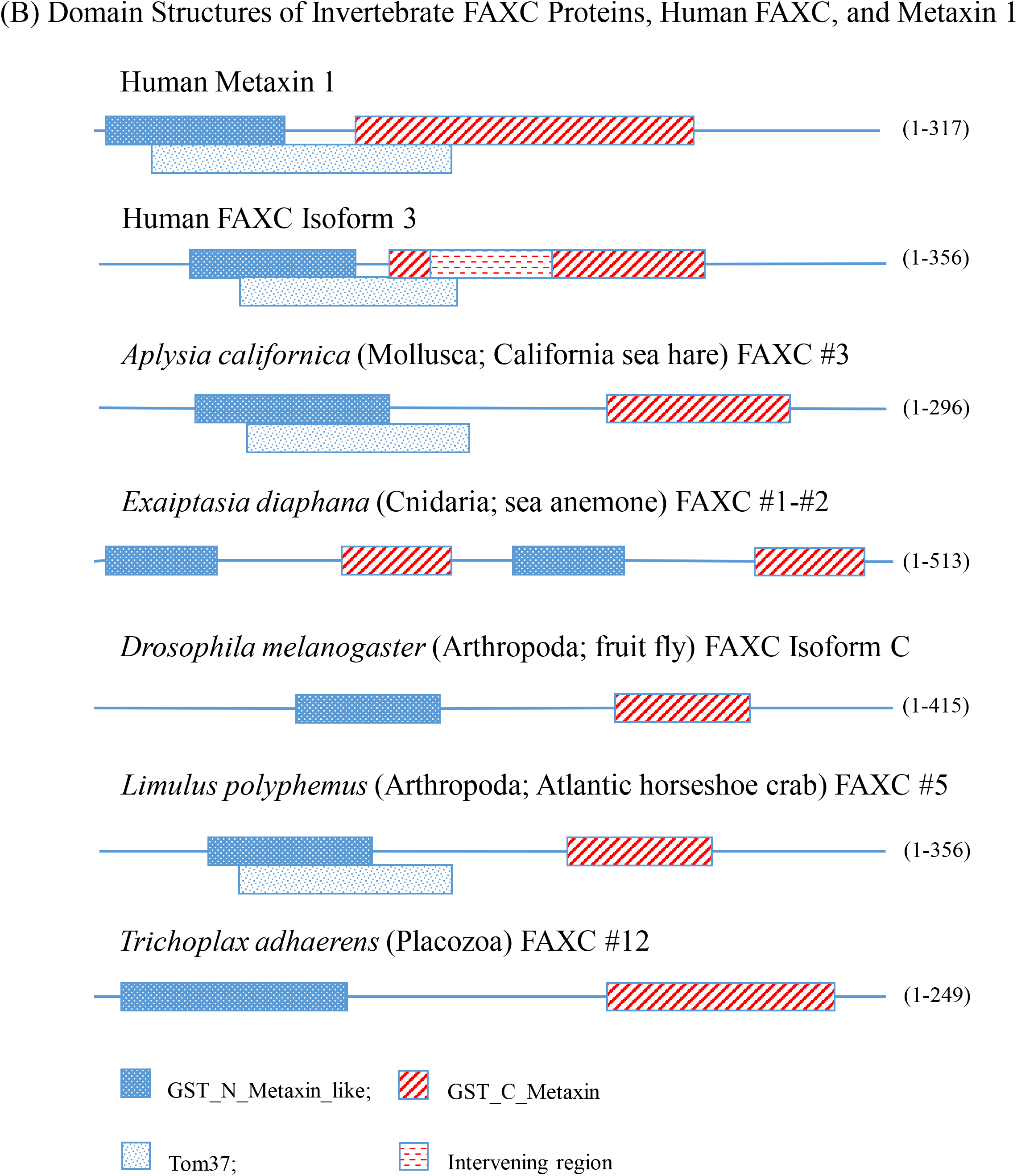
Major protein domains of FAXC and metaxin proteins of vertebrates and invertebrates. (A) The characteristic GST_N_Metaxin and GST_C_Metaxin protein domains of representative vertebrate FAXCs and metaxins are shown. The conserved Tom37 domain is also included in the figure. The three domains are present in human metaxins 1, 2, and 3 and are also major domains of the human FAXC protein. Isoform 1 of human FAXC is shown, but the domains of isoforms 2 and 3 are identical. The isoforms differ in the length of the N-terminal sequences. Isoform 1 (409 aa) has an extra 121 N-terminal amino acids compared to isoform 2 (288 aa), and an extra 53 compared to isoform 3 (356 aa). In addition to the human FAXC isoform 1 domains, the FAXC domains of mouse, *Danio rerio* (zebrafish), and *Xenopus laevis* (frog) are included in the figure. They all have the same GST_N_Metaxin, GST_C_Metaxin, and Tom37 domain structure. However, human, mouse, and *Xenopus* have broken GST_C_Metaxin domains, with an inserted, intervening protein sequence. Figure 1B includes the conserved protein domains of representative invertebrate FAXCs, as well as human metaxin 1 and human FAXC isoform 3. For all examples, the GST_N_Metaxin and GST_C_Metaxin domains are the major, characteristic protein domains. However, the Tom37 domain is absent from some invertebrate FAXCs. A variety of invertebrate phyla have species that possess FAXCs with the GST_N_Metaxin and GST_C_Metaxin domains. These include Mollusca (example in the figure: the sea hare *Aplysia californica*), Cnidaria (the sea anemone *Exaiptasia diaphana*), Arthropoda (the fruit fly *Drosophila melanogaster*; the horseshoe crab *Limulus polyphemus*), and Placozoa (*Trichoplax adhaerens*). For *Exaiptasia*, the genes for FAXC #1 and FAXC #2 are contiguous.

**Figure 2.**
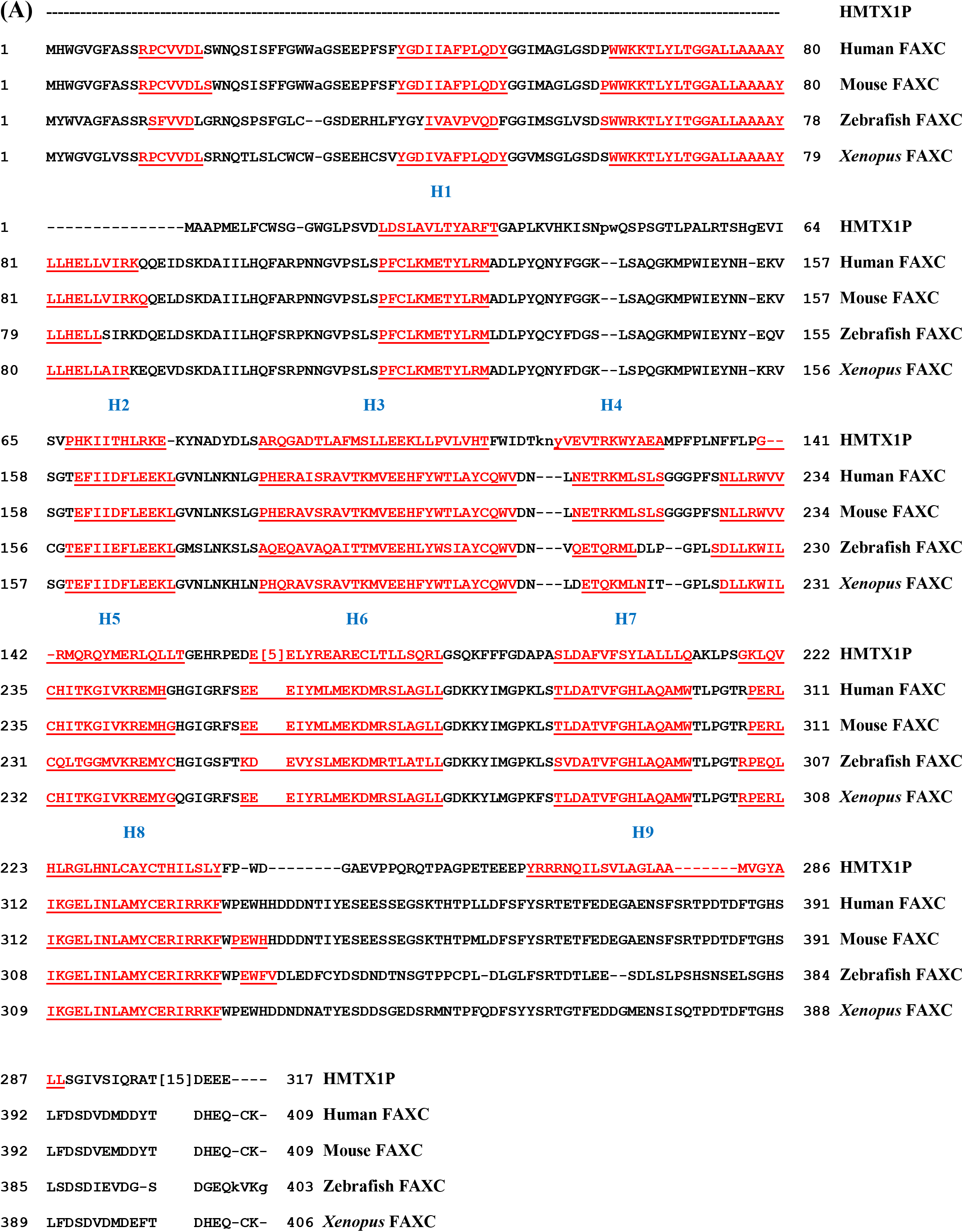
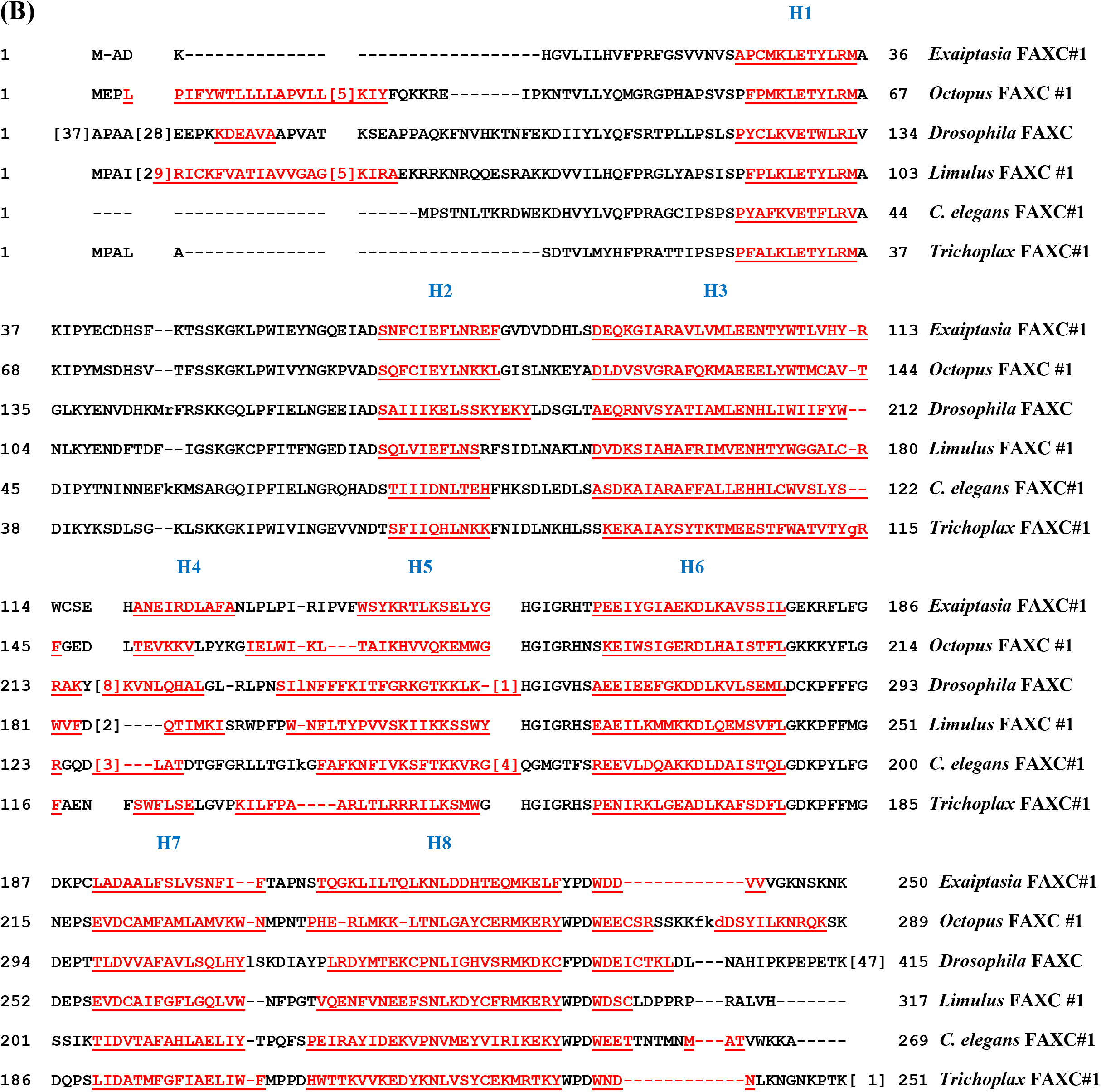
Alpha-helical secondary structures of FAXC proteins of vertebrates and invertebrates. (A) As shown in the figure, multiple sequence alignments demonstrate that α-helical segments dominate the predicted FAXC protein secondary structures of representative vertebrates. The α-helical segments are highlighted in red and underlined, and are labeled H1 through H8 based on the α-helices of human metaxin 1 (HMTX1P). The FAXCs in the figure include human FAXC isoform 1, mouse (*mus musculus*) FAXC, and the FAXCs of zebrafish (*Danio rerio*) chromosome 16 and *Xenopus laevis* chromosome 5L. The figure shows that the patterns of α-helices H1 – H8 are highly conserved among the vertebrate FAXC proteins. Also, the patterns of helices for the FAXCs are seen to be identical to the pattern for human metaxin 1. In contrast, there are few β-strand segments in the FAXC or metaxin 1 proteins. The FAXCs have extra helices that precede H1 at the N-terminal ends of the proteins. In addition, human metaxin 1 has an extra C-terminal helix H9. (B) Invertebrates of a variety of phyla also have FAXC proteins with secondary structures dominated by α-helices, and with a pattern of helices, H1 through H8, that is highly conserved. The invertebrates in the figure represent phyla that include Cnidaria (*Exaiptasia diaphana*), Mollusca (*Octopus bimaculoides*), Arthropoda (*Drosophila melanogaster*, *Limulus polyphemus*), Nematoda (*C. elegans*), and Placozoa (*Trichoplax adhaerens*). FAXC #1 is shown for all of the examples, which have multiple FAXCs, except for the *Drosophila melanogaster* FAXC, which is isoform C. Conserved α-helices and similar spacings between helices are therefore the rule for both vertebrates and invertebrates.

**Figure 3.**
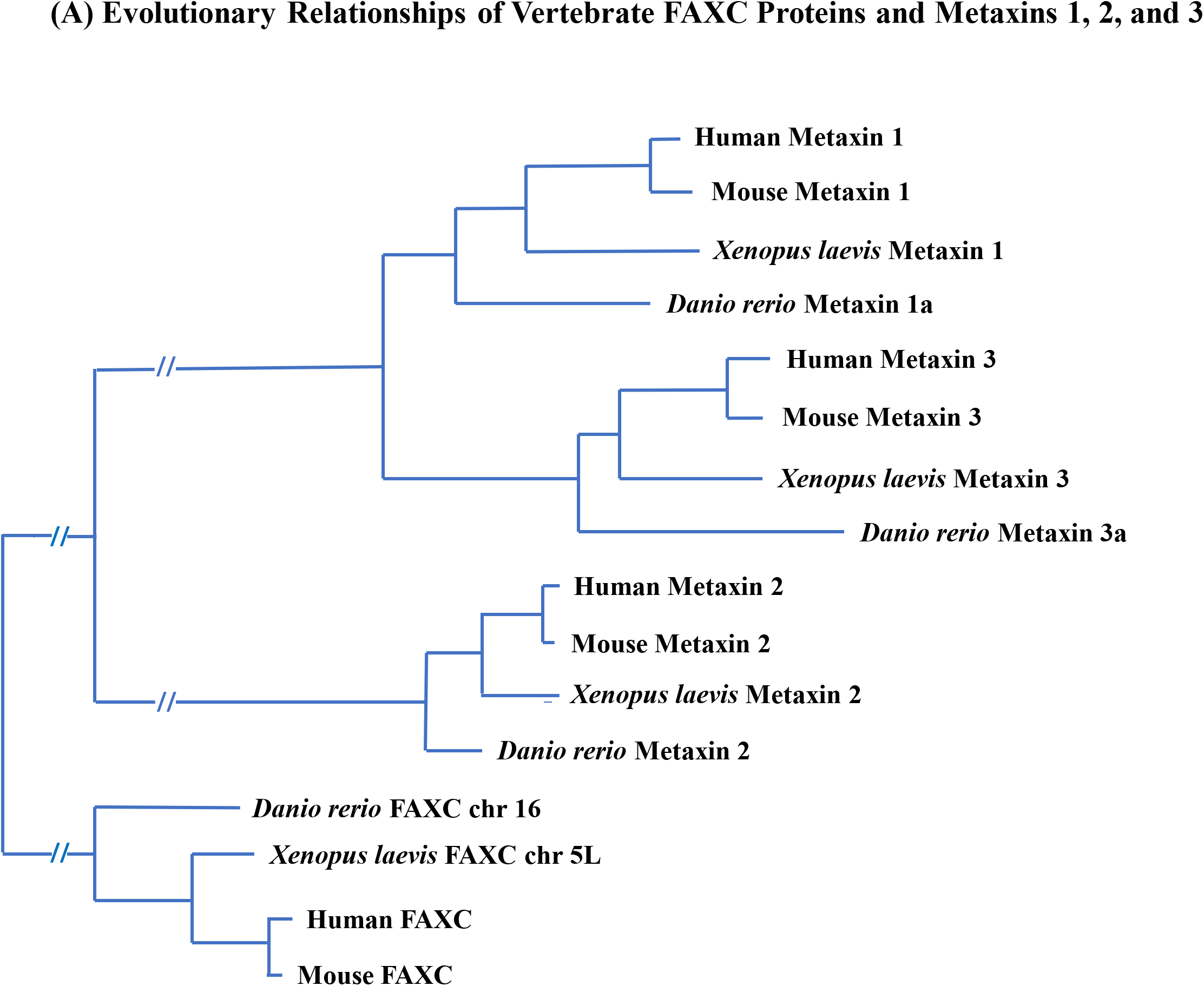
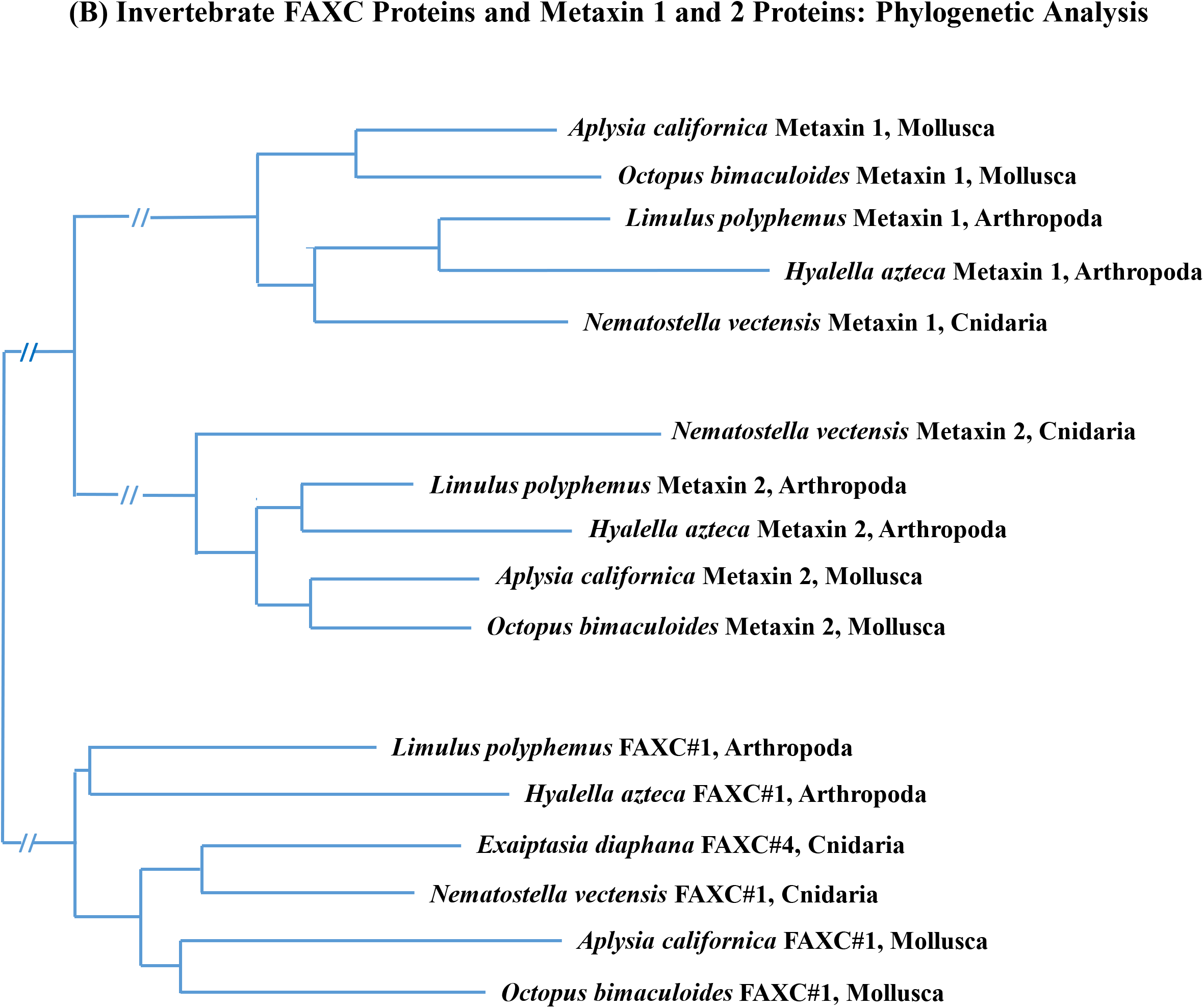
Evolutionary relationships of FAXC and metaxin proteins of vertebrates and invertebrates. (A) Phylogenetic analysis of vertebrate FAXCs and metaxins: human, mouse, *Xenopus laevis*, and *Danio rerio*. The phylogenetic tree shows separate groups of metaxin 1, metaxin 2, metaxin 3, and FAXC proteins. Within each group, human and mouse are most related, then *Xenopus laevis*, and then *Danio rerio*. For the metaxins, the metaxin 1 and 3 proteins have greater similarity compared to the metaxin 2 proteins. Notably, the FAXC proteins are seen to be phylogenetically related to the vertebrate metaxins, a result compatible with the conserved protein domains and α-helical structures of both the FAXCs and metaxins. In the figure, the extent of evolutionary change is proportional to the horizontal lengths of the branches. (B) Phylogenetic relationships of invertebrate FAXC and metaxin proteins. The metaxin 1, metaxin 2, and FAXC proteins of invertebrates of three phyla that possess FAXCs are included in the figure. These phyla are Mollusca (*Aplysia californica*, *Octopus bimaculoides*), Arthropoda (*Limulus polyphemus*, *Hyalella azteca*), and Cnidaria (*Nematostella vectensis*, *Exaiptasia diaphana*). The proteins form three separate groups: metaxin 1, metaxin 2, and FAXC. Within each group, the metaxins or FAXCs of species of the same phylum are most closely related, for example the molluscs *A. californica* and *O. bimaculoides*. Significantly, the figure demonstrates that the three groups of invertebrate proteins – metaxins 1 and 2 and FAXCs – are related by evolution.

**Figure 4.**
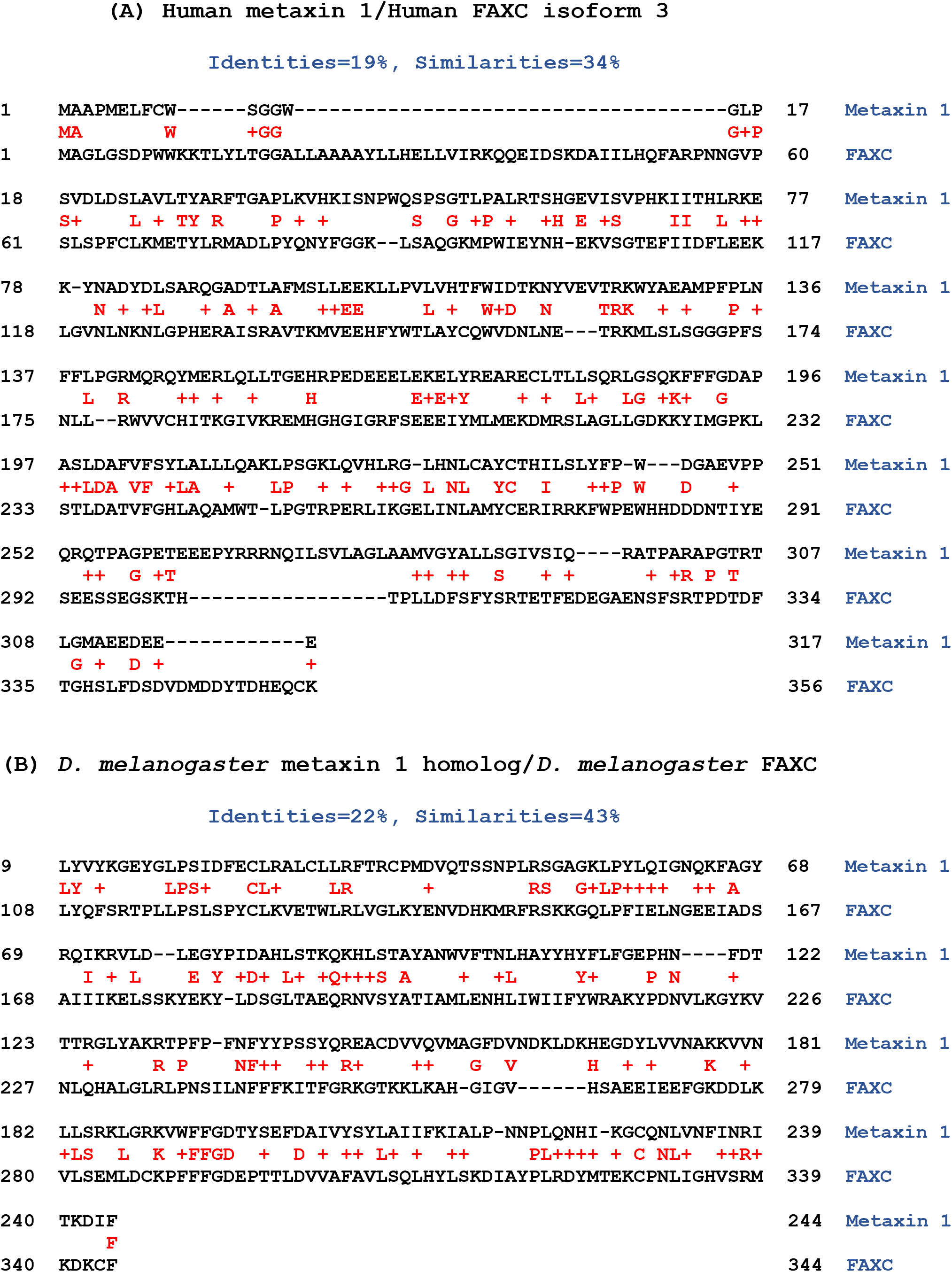
Alignment of FAXC and metaxin amino acid sequences. In Figure 4A, the amino acid sequence of human metaxin 1 is aligned with the sequence of human FAXC isoform 3. Shown between the sequences in red are the identical amino acids, with the similar amino acids indicated by + signs. Only 19% identical amino acids and 34% similar amino acids are found using NCBI Global Align. With NCBI Align Two Sequences, which only includes the regions of greatest homology, there are 26% identities. Similar results are found for other vertebrates. These low percentages indicate that FAXC and metaxin proteins represent different categories of proteins, even though they share structural features such as the same protein domains and α-helical segments. This conclusion is confirmed by sequence alignments of invertebrate FAXC and metaxin proteins (Figure 4B). The example in the figure includes the amino acid sequence of the *Drosophila melanogaster* metaxin 1 homolog (CG9393) aligned with the sequence of *D*. *melanogaster* FAXC isoform C. There are 22% identical amino acids with Align Two Sequences, as shown in the figure, and 14 % with Global Align. Invertebrates of different phyla show comparable results.

**Figure 5.**
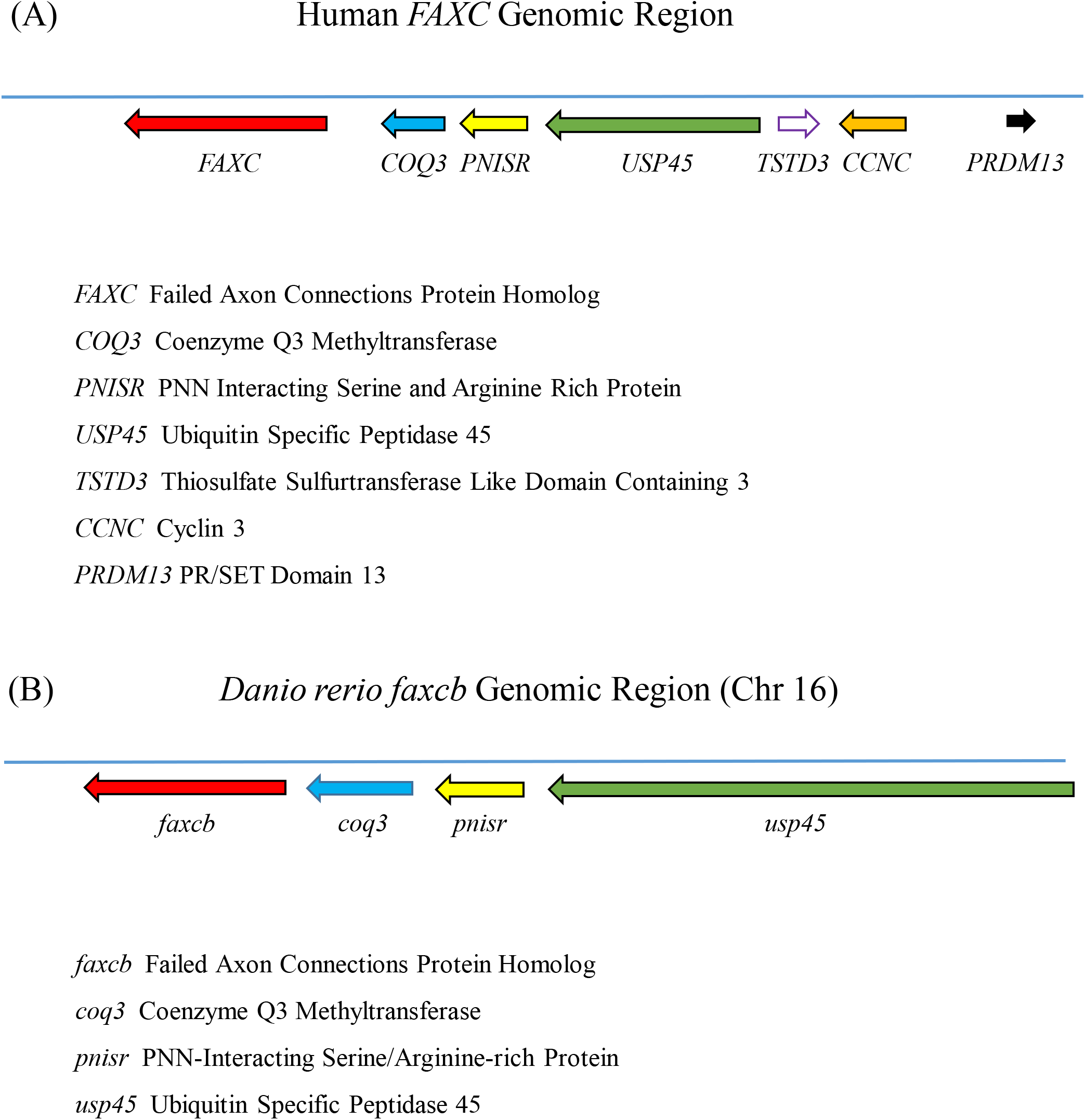
Genomic regions of vertebrate FAXC genes. (A) Neighboring genes of the human *FAXC* gene. The genes that are adjacent to the *FAXC* gene are listed on the figure and are in the order *FAXC* – *COQ3* – *PNISR* – *USP45* – *TSTD3* – *CCNC* – *PRDM13*. The neighboring genes of other vertebrates such as *Danio rerio*, shown in (B), and *Xenopus laevis* are similar and have a similar order. The genomic region of the human metaxin 1 gene (*MTX1*) at cytogenetic location 1q22 has a different set of genes compared to the human *FAXC* gene at 6q16.2. (B) Neighboring genes of the *Danio rerio faxcb* gene. The zebrafish genome has two *faxc* genes due to genome duplication: *faxca* on chromosome 4 and *faxcb* on chromosome 16. The *faxcb gene*, which is shown in the figure, has the same neighboring genes as the human *FAXC* gene in (A). They are also in the same order: *faxcb* – *coq3* – *pnisr* – *usp45*. The *faxca* genomic region on chromosome 4, however, shows neighboring genes that are largely different.

**Figure 6.**
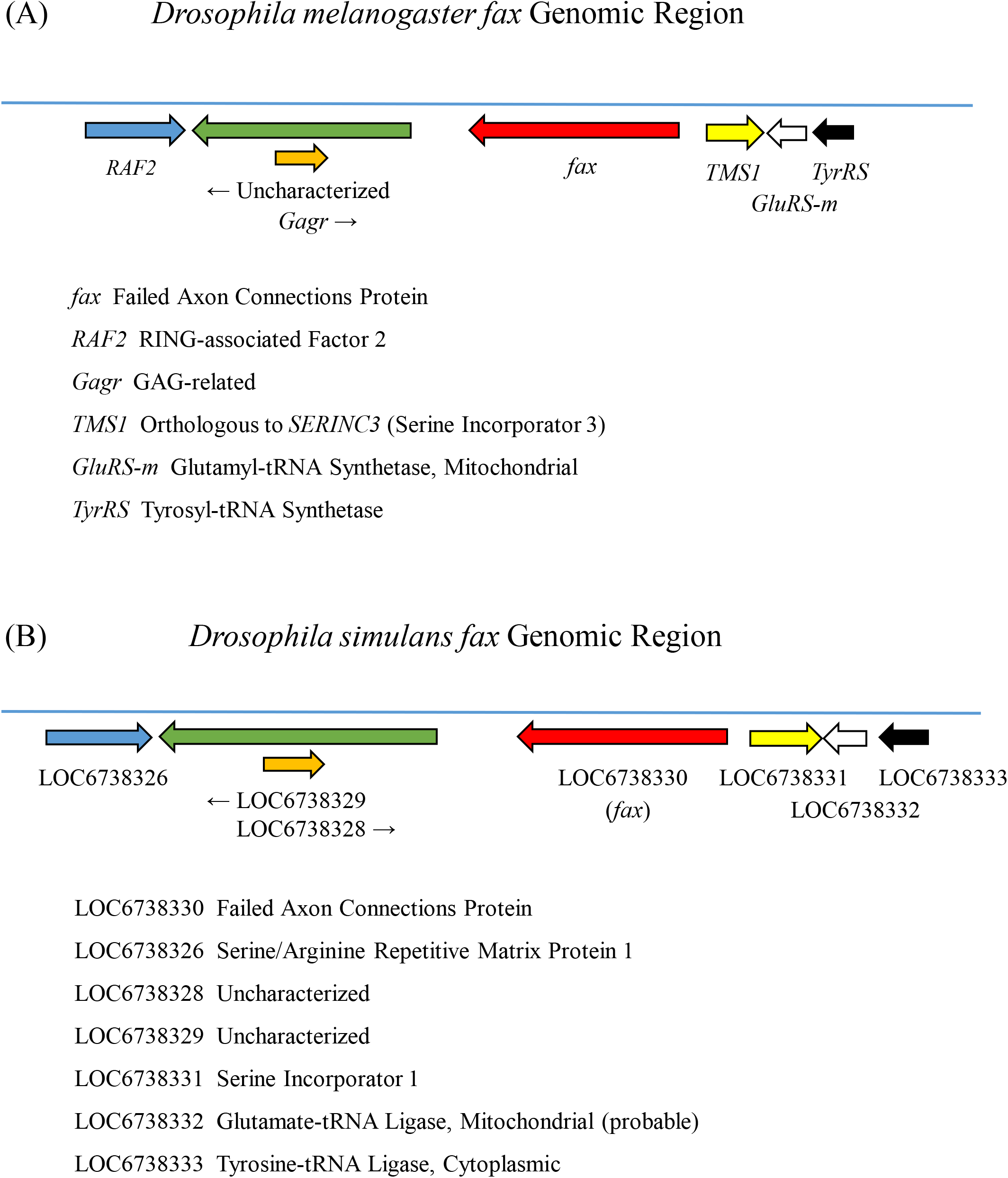
Genomic regions of invertebrate FAXC genes. (A) Adjacent genes of the *Drosophila melanogaster fax* gene. As shown in the figure, the genes next to the *D*. *melanogaster fax* gene are different than the genes next to the human and *Danio rerio* FAXC genes (Figure 5), and also the FAXC genes of *Xenopus laevis* and other vertebrates. The *Drosophila* genes are listed on the figure and are in the order *RAF2* – *Uncharacterized* – *Gagr* – *fax* – *TMS1* – *GluRS-m* – *TyrRS*. The results with additional invertebrate and vertebrate FAXCs (not shown) strengthen the conclusion that invertebrate FAXC genes have different neighboring genes than vertebrate FAXCs. (B) The *Drosophila simulans fax* gene region. The genes adjacent to the *fax* gene and their order are the same for *D. melanogaster* and *D. simulans*, a related fruit fly species of the melanogaster subgroup. This is in accord with the expectation that related invertebrate species are likely to have similar *fax* genomic regions.

**Figure 7.**
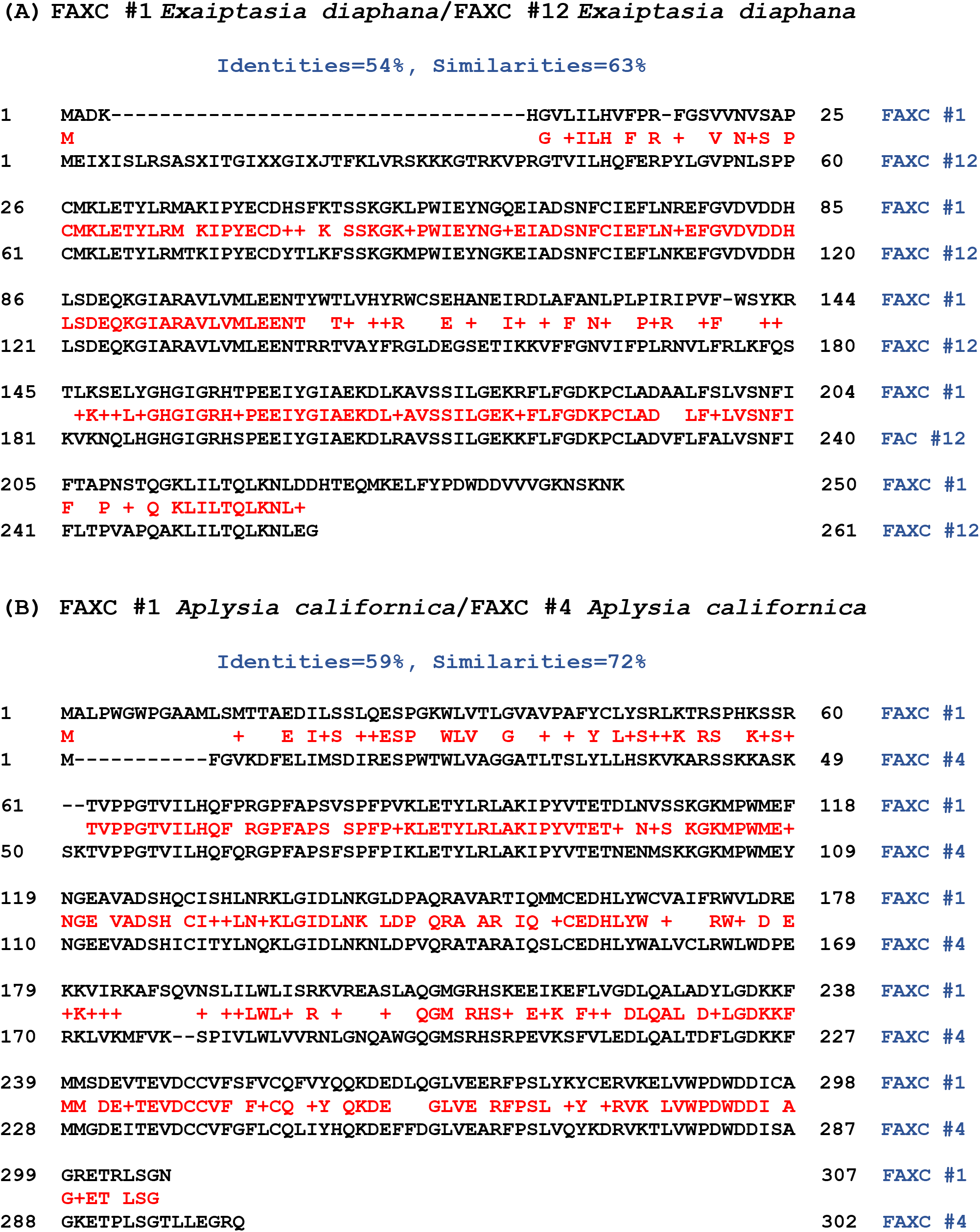
Multiple FAXC proteins of invertebrates: amino acid sequence alignments. In (A), FAXC #1 of the sea anemone *Exaiptasia diaphana* (Cnidaria) is aligned with FAXC #12 of *Exaiptasia*. The letters and “+” signs between the sequences and highlighted in red denote the amino acid identities and similarities, respectively. The percent identities (54% with NCBI Global Align as shown in the figure, 69% with Align Two Sequences) indicate that the multiple FAXCs share a high level of homology. A second example of an invertebrate with multiple predicted FAXC proteins is shown in Figure 7B. The figure compares FAXC #1 with FAXC #4 of the California sea hare *Aplysia californica* (Mollusca). As in (A), a high percentage of identical amino acids is found (59% with Global Align as included in the figure, 63% with Align Two Sequences). Similar results, showing high levels of identical amino acids for multiple FAXCs, are found for other invertebrates with more than one FAXC, such as *Branchiostoma floridae* (8 FAXCs) and *Trichoplax adhaerens* (12 FAXCs).

**Figure 8.**
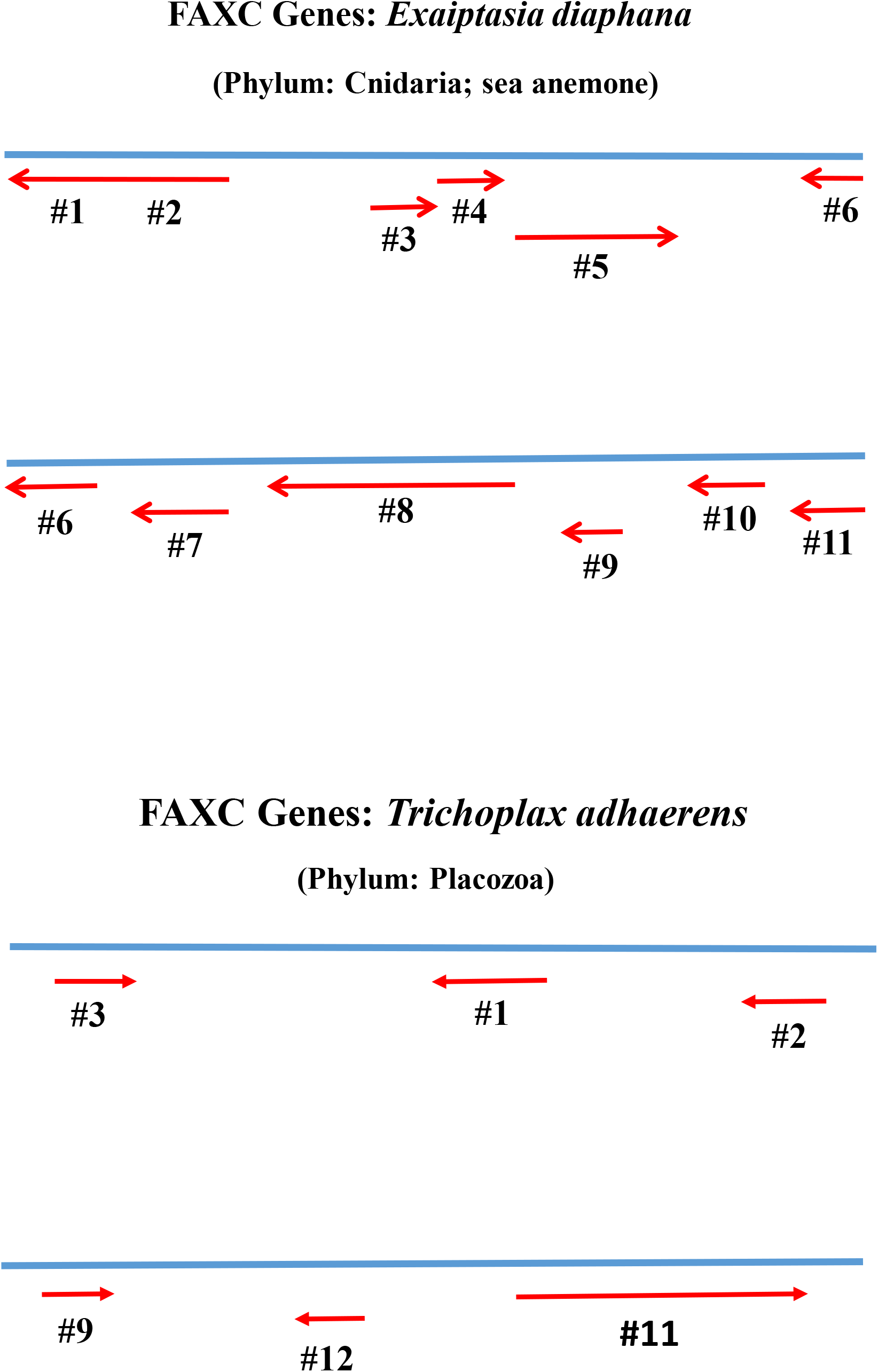
Multiple FAXC genes in invertebrates: genomic regions. The figure shows two examples of genomic regions of invertebrates with multiple FAXC genes that are in close proximity. At the top of the figure, a segment of the *Exaiptasia diaphana* genome is included with 11 FAXC genes (#1 - #11) that are clustered in the same genomic region. At least 13 FAXC genes exist in *Exaiptasia*, and most (11 of 13) are therefore in the same localized region of the genome. At the bottom of the figure, a segment of the *Trichoplax adhaerens* genome is included which has 6 of the 12 FAXC genes that have been identified. The other 6 FAXC genes are distributed throughout different regions of the *Trichoplax* genome.

## 3. RESULTS AND DISCUSSION

### 3.0. FAXC Proteins: Widely Occurring Proteins with Structural Similarity to Metaxins

As described in this report, searches of protein databases have revealed that FAXC proteins are widely distributed proteins that have features in common with the metaxins and can be considered metaxin-like. These features include the characteristic GST_N_Metaxin and GST_C_Metaxin protein domains of metaxins and a pattern of α-helical segments resembling that of metaxin 1 and other metaxins.

While FAXC proteins were originally identified in *Drosophila*, homologous proteins with metaxin-like structural features are shown in this report to exist in a wide variety of species. In addition to vertebrates and invertebrates with FAXC homologs, bacteria and fungi also have been found to possess FAXC-like genes (K.W. Adolph, unpublished). For invertebrates, the phyla that have the largest number of species with FAXC proteins are Mollusca, Arthropoda including Insecta and Arachnida, Cnidaria, and Placozoa. Examples include octopus and snail (Mollusca), lobster and water flea (Arthropoda), bee and ant (Insecta), scorpion and spider (Arachnida), coral and sea anemone (Cnidaria), and *Trichoplax* (Placozoa).

Not all invertebrates appear to possess FAXC genes. Phyla for which FAXC genes have not yet been detected include Echinodermata (example: sea urchins) and Porifera (sponges). Likewise, FAXC-like genes have not been observed in flowering plants and archaea. However, the list of organisms with FAXC genes may lengthen as new genome sequences are added to the NCBI and other databases.

The possession of multiple FAXC genes is a notable feature of invertebrates. *Exaiptasia diaphana* (sea anemone; Cnidaria) has at least 13 FAXC genes, *Trichoplax adhaerens* (Placozoa) at least 12, and *Branchiostoma floridae* (Florida lancelet; Chordata) at least 10. Numerous other examples in different invertebrate phyla also have multiple FAXC proteins.

### 3.1. Conserved Protein Domains of FAXC Proteins of Vertebrates and Invertebrates

A structural feature of proteins that is important in identifying the proteins as metaxins or metaxin-like is the presence of conserved metaxin protein domains. These are the GST_N_Metaxin and GST_C_Metaxin domains. They are found in vertebrate metaxins 1, 2, and 3, invertebrate metaxins 1 and 2, and the metaxin-like proteins of plants, protists, fungi, and bacteria. And they are a major structural feature of FAXC proteins, as demonstrated in this paper.

Figure 1A includes the major conserved protein domains of the FAX proteins of a selection of vertebrates, along with the major protein domains of human metaxins 1, 2, and 3. The metaxins in the figure possess the characteristic GST_N_Metaxin and GST_C_Metaxin domains, as well as Tom37 domains. Importantly, the FAXC proteins in Figure 1A also have the same major conserved domains. Along with sharing the same domains, FAXC proteins and metaxins have other structural similarities, in particular similar patterns of α-helical segments, as discussed in Section 3.2. Another criterion for being a metaxin is the possession of a high degree of protein sequence homology with the metaxins. But amino acid sequence alignments, such as in Figure 4A, reveal only low percentages of sequence identity in comparing vertebrate FAXC proteins and metaxins. Taking into account the criterion of sequence homology, FAXC proteins are not identical to the metaxins, but do have metaxin-like structural features.

The four examples of vertebrate FAXCs in Figure 1A (human, mouse, zebrafish, and *Xenopus*) have metaxin 1 domains rather than metaxin 2 domains. This is generally true of vertebrate FAXCs. However, amino acid sequence alignments show that vertebrate FAXCs are about equally similar to metaxins 1, 2, and 3, and do not have substantially greater similarity to metaxin 1.

In addition to the GST_N_Metaxin and GST_C_Metaxin domains, the Tom37 conserved protein domain may also be present in FAXC proteins. The four vertebrate examples in Figure 1A are seen to possess the Tom37 domain. The Tom37 domain was initially identified in the Sam37 protein of the yeast *Saccharomyces cerevisiae*. The Sam37 protein is a component of the SAM complex of the outer mitochondrial membrane of *Saccharomyces cerevisiae* and related species, and is involved in the uptake of proteins into mitochondria.

Both zebrafish and *Xenopus* have two FAXC genes due to genome duplication: for zebrafish, *faxca* (chromosome 4) and *faxcb* (chr 16), and for *Xenopus, faxc.L* (chr 5L) and *faxc.S* (chr 5S). The two genes for both zebrafish and *Xenopus* encode proteins with similar domain structures. Figure 1A includes the zebrafish FAXC protein encoded by *faxcb* on chromosome 16 and the *Xenopus* FAXC protein encoded by *faxc.L* on chromosome 5L.

Figure 1B shows the domain structures of representative invertebrate FAXCs. Also included are human metaxin 1 and human FAXC isoform 3. Like the FAXCs of vertebrates, invertebrate FAXCs are characterized by GST_N_Metaxin and GST_C_Metaxin domains. For invertebrates, the Tom37 domain is present in some but not all FAXC proteins. The Tom37 domain is absent in the insect example (*Drosophila melanogaster*) and in *Trichoplax*. FAXCs of a variety of invertebrates are shown in the figure, including FAXCs of Mollusca, Cnidaria, Arthropoda, and Placozoa.

The invertebrates in Figure 1B are important model organisms of biological research or have economic or medical significance. These include the California sea hare *Aplysia californica*, the sea anemone *Exaiptasia diaphana*, the fruit fly *Drosophila melanogaster*, the Atlantic horseshoe crab *Limulus polyphemus*, and the placozoan *Trichoplax adhaerens*. Some additional information and references regarding the selected invertebrates are given in the Introduction. The example of *Exaiptasia diaphana* shows two FAXC genes (#1 and #2) that are contiguous (see also Figure 8). *Exaiptasia*, like a number of other invertebrates, has multiple FAXC genes, in this case at least 13.

### 3.2. Alpha-Helical Structures of Vertebrate and Invertebrate FAXC Proteins

Vertebrate and invertebrate FAXCs have patterns of α-helical segments similar to the metaxins, in addition to having the same major protein domains. This is illustrated in Figure 2A for vertebrate FAXCs. Human, mouse, zebrafish, and *Xenopus* FAXCs have patterns of secondary structure dominated by α-helices. In particular, eight α-helices, H1 through H8, that characterize the structure of human metaxin 1 and other metaxins are also the major feature of vertebrate FAXC secondary structures. In addition, similar spacings between the α-helical segments are found. Human metaxin 1 has an extra C-terminal helix H9, while the FAXCs have extra N-terminal helices.

The secondary structures of vertebrate FAXCs and metaxins are highly conserved, as shown by the examples in Figure 2A, even though only low percentages of amino acid identities are revealed in aligning pairs of FAXC and metaxin sequences. For instance, human FAXC isoform 1 and human metaxin 1 have only 17% identical amino acids (using NCBI Global Align), while zebrafish Faxcb and human metaxin 1 also have 17% identical amino acids. But all these sequences have the conserved α-helical segments H1 – H8.

Invertebrate FAXC secondary structures are also characterized by the presence of helices H1 through H8 (Figure 2B). The figure includes the FAXC proteins of invertebrates of major invertebrate phyla: Cnidaria (*Exaiptasia*), Mollusca (*Octopus*), Arthropoda (*Drosophila*, *Limulus*), Nematoda (*C. elegans*), and Placozoa (*Trichoplax*).

As the examples in Figure 2B demonstrate, invertebrate FAXCs, like metaxins 1, 2, and 3, share a high level of conservation of α-helices H1 – H8. However, similar to the vertebrate FAXCs, they display only low percentages of amino acid identities aligned with metaxin proteins. For instance, human metaxin 1 and *Exaiptasia diaphana* FAXC #1 have only 21% identities, and human metaxin 1 and *Octopus bimaculoides* FAXC #1 have only 18%.

A transmembrane α-helix, predicted from the amino acid sequence (Krogh et al., 2001; Kall et al., 2007), is a characteristic feature of vertebrate metaxin 1 proteins, but not metaxin 2 or metaxin 3 proteins. The α-helical segment is near the C-terminus of metaxin 1 proteins, for example between residues 272 and 294 of human metaxin 1 (317 amino acids). It may serve to anchor metaxin 1 to the outer mitochondrial membrane. The C-terminal transmembrane α-helix is also present in metaxins or metaxin-like proteins of invertebrates, plants, protists, and fungi, but not bacteria. However, transmembrane helices were not found for FAXC proteins of vertebrates or invertebrates. The vertebrates included human, mouse, zebrafish, and *Xenopus*. The invertebrates included *Exaiptasia*, *Octopus*, *Drosophila*, *Limulus*, *C. elegans*, *Trichoplax*, *Aplysia*, *Nematostella*, and *Hyalella*.

### 3.3. Phylogenetic Analysis of Vertebrate and Invertebrate FAXCs and Metaxins

Evolutionary relationships between vertebrate FAXC proteins and vertebrate metaxins are shown in Figure 3A. Human, mouse, *Xenopus*, and zebrafish proteins are included in the figure. The FAXC proteins and vertebrate metaxins 1, 2, and 3 all form separate groups. In each group, human and mouse are most closely related, followed by *Xenopus*, and then zebrafish. For the vertebrate FAXCs, phylogenetic results such as in Figure 3A indicate that the FAXCs are related by evolution to the metaxins. The phylogenetic results are in keeping with the conserved protein domain structures and α-helical secondary structures of FAXCs and metaxins. Among the metaxins, the metaxin 1 and metaxin 3 proteins are most closely related, while the metaxin 2 proteins are somewhat less related. These results are compatible with amino acid sequence alignments. In particular, human metaxins 1 and 3 share 45% identical amino acids, but human metaxins 1 and 2 only 22%.

Figure 3B shows the phylogenetic relationships of invertebrate FAXC proteins and metaxins. Invertebrates have metaxin 1 and metaxin 2 proteins like vertebrates, and metaxins 1 and 2 of a selection of invertebrates, along with FAXC proteins, are included in the figure. The FAXC proteins of invertebrates of the same phylum, such as *Aplysia* and *Octopus* FAXCs (both Mollusca), are seen to be most closely related, in keeping with their taxonomic classification. Metaxin proteins of the same phylum are also most closely related. Other invertebrate phyla in the figure are Arthropoda (*Limulus* and *Hyalella*) and Cnidaria (*Nematostella* and *Exaiptasia*). The three types of proteins (metaxin 1, metaxin 2, and FAXC) form three distinct groups. But the groups are related by evolution, as with the vertebrates.

### 3.4. Amino Acid Sequence Alignments of FAXC and Metaxin Proteins of Vertebrates and Invertebrates

FAXC proteins have metaxin-like features, as discussed in the previous sections. The similarities of the domain structures and α-helical segments of vertebrate and invertebrate FAXCs and metaxins, along with the phylogenetic results, clearly demonstrate that FAXC proteins are metaxin-like. But amino acid sequence alignments show that, while FAXCs have metaxin-like features, they cannot simply be categorized as metaxins. As an example, Figure 4A includes the alignment of human metaxin 1 (317 amino acids) and human FAXC isoform 3 (356 aa). Only 19% amino acid identities are found using NCBI Global Align. Using Align Two Sequences gives 26% identities.

Other vertebrates also reveal low percentages of identical amino acids in aligning FAXC and metaxin protein sequences. For the zebrafish *Danio rerio*, alignment of the Faxcb protein (403 amino acids) encoded by *faxcb* with the zebrafish metaxin 1a protein (Mtx1a; 319 aa) encoded by *mtx1a* shows only 19% identical amino acids. Alignment of Faxcb with metaxin 1b (Mtx1b; 317 aa) encoded by *mtx1b* also gives 19% identities. Besides having two FAXC genes, *faxca* on chromosome 4 and *faxcb* on chromosome 16, *Danio rerio* has two metaxin 1 genes, *mtx1a* on chromosome 16 and *mtx1b* on chromosome 19. For the frog *Xenopus laevis*, just 18% identical amino acids are found in aligning the Faxc.S protein encoded by *faxc.S* with *Xenopus* metaxin 1 S encoded by *mtx1.S*. Likewise, alignment of *Xenopus* Faxc.L encoded by *faxc.L* with *Xenopus* metaxin 1 S also reveals only 18% identities.

Figure 4B includes an example of the alignment of an invertebrate metaxin 1 and FAXC. The invertebrate is *Drosophila melanogaster*. The *Drosophila* metaxin 1 homolog (CG9393; 315 amino acids) and FAXC isoform C (415 aa) have 22% identical amino acids with Align Two Sequences, and only 14% with Global Align. The low percentages of identical amino acids in aligning metaxin 1 and FAXC proteins of the human and *Drosophila* examples indicate that metaxins and FAXCs are separate categories of proteins, but with structural features in common.

The results with metaxin 2 homologs and FAXC proteins are comparable. Human metaxin 2 (263 aa) and human FAXC isoform 3 (356 aa) have 19% identical amino acids with Global Align, and 25% with Align Two Sequences. *Drosophila melanogaster* has two metaxin 2 homologs: CG8004 (269 aa) and CG5662 (292 aa). CG8004 and *Drosophila* FAXC isoform C have 15% identities (Global Align), while CG5662 and FAXC isoform C have 17% identities (Global Align). These results strengthen the conclusion that FAXCs have metaxin-like features, but are in a different category than the metaxins.

### 3.5. Genomic Regions of FAXC Genes: Neighboring Genes

The genes adjacent to the human *FAXC* gene and the *Danio rerio faxcb* gene are shown in Figure 5A,B as examples of the neighboring genes of vertebrate FAXCs. The figure demonstrates that the genes adjacent to vertebrate FAXC genes are largely conserved for these examples. The same order of genes is found for both genomic regions: *FAXC* – *COQ3* – *PNISR* – *USP45* (for human *FAXC* at cytogenetic location 6q16.2) and *faxcb* – *coq3* – *pnisr* – *usp45* (for zebrafish *faxcb* on chromosome 16). The gene arrangement with other vertebrates is generally similar to the human and zebrafish arrangements. For example, the *Xenopus faxc.L* gene on chromosome 5L has neighboring genes that are the same as the genes next to the human *FAXC* gene and the zebrafish *faxcb* gene. However, the zebrafish *faxca* genomic region, on chromosome 4, has different neighboring genes: *faxca* – *ccnc* (cyclin c) – *ngs* (notochord granular surface) –*pgm3* (phosphoglucomutase 3).

The genes adjacent to FAXC genes differ from the genes adjacent to metaxin genes. For example, the human metaxin 1 gene at cytogenetic location 1q22 is between the genes for thrombospondin 3 (*THBS3*) and a glucosylceramidase beta 1 pseudogene (*psGBA1*). The human metaxin 3 gene at 5q14.1 is between the thrombospondin 4 gene (*THBS4*) and the cardiomyopathy associated 5 gene (*CYMA5*). And the metaxin 2 gene is next to a group of homeobox genes.

For invertebrates, the FAXC genomic regions are different than the vertebrate FAXC regions. This can be seen in Figure 6A,B, which includes the genes that are next to the *fax* genes of *Drosophila melanogaster* and *Drosophila simulans*. For *D. melanogaster*, the genes are in the order *RAF2* – *Uncharacterized* – *Gagr* – *fax* – *TMS1* – *GluRS-m* – *TyrRS*. A similar gene order is found for *D*. *simulans*. The FAXC genomic regions for vertebrates (Figure 5) and invertebrates (Figure 6) are therefore very different. In addition, comparing Figure 6A with 6B confirms that the gene orders for the invertebrate FAXC genomic regions of closely related but distinct species are typically identical or very similar. In this case, *D. melanogaster* and *D. simulans* are closely related and are both in the melanogaster species taxonomic subgroup.

### 3.6. Multiple FAXC Genes in Invertebrates

A characteristic of the metaxin-like proteins of fungi such as *Schizosaccharomyces pombe*, *Aspergillus nidulans*, and *Penicillium rubens* is the presence of multiple genes, MTXa and MTXb (Adolph, 2022). Multiple metaxin genes are also found with vertebrates. For instance, the zebrafish *Danio rerio* has two metaxin 1 genes, *mtx1a* and *mtx1b*. But, compared to the metaxins and metaxin-like proteins, an even greater multiplicity of FAXC genes is found for invertebrates. The sea anemone *Exaiptasia diaphana*, for example, has at least 13 FAXC genes. The placozoan *Trichoplax adhaerens* has at least 12. The lancelet *Branchiostoma floridae* has at least 10. And the coral *Stylophora pistillata*, 9. Multiple FAXC genes are especially common among invertebrate phyla that include Cnidaria, Mollusca, Arthropoda, Placozoa, Chordata, and Nematoda. Vertebrates, in contrast, appear to have primarily a single FAXC gene. However, there are exceptions. These include *Danio rerio* which has two FAXC genes, *faxca* and *faxcb*, due to genome duplication in fish.

Figure 7A and Figure 7B show examples of amino acid sequence alignments of pairs of FAXC protein sequences of invertebrates with multiple FAXCs. High percentages of sequence identities and similarities are found for both examples. In (A), FAXC #1 of the sea anemone *Exaiptasia diaphana* (Cnidaria) is aligned with FAXC #12. The identities are 54% using NCBI Global Align (69% using Align Two Sequences). In (B), FAXC #1 of *Aplysia californica* (Mollusca), the California sea hare, is aligned with FAXC #4, and reveals 59% identities with Global Align (63% with Align Two Sequences). Comparable high levels of sequence identities are found with different pairs of FAXC proteins of *Exaiptasia* and *Aplysia*.

An additional example is the alignment of FAXC #1 of the Florida lancelet *Branchiostoma floridae* (Chordata) with FAXC #8. The amino acid identities are 49% using Global Align and 51% using Align Two Sequences. As a final example, *Trichoplax adhaerens* (Placozoa) FAXC #1 and FAXC #12 have 38% identities with Global Align and 41% with Align Two Sequences.

The high percentages of amino acid identities in aligning the sequences of different FAXC proteins of invertebrates that have multiple FAXC genes suggest that the multiple FAXC genes encode proteins with conserved structures. However, since little is known about the expression of FAXC genes or the functions of FAXC proteins, the significance of the structures of multiple FAXC proteins remains to be determined.

The multiple FAXC genes may be distributed in different regions of the genome, or may be arranged as a cluster of genes. Figure 8 (Top) displays a cluster of multiple FAXC genes of *Exaiptasia diaphana*. The cluster includes 11 of the 13 or more FAXC genes, labelled #1 through #11, of this cnidarian. No other genes are between the FAXC genes. Eight of the genes are oriented right to left on one DNA strand, and 3 left to right on the other strand. Figure 8 (Bottom) shows 6 of the 12 FAXC genes of *Trichoplax adhaerens*. Three are oriented right to left on one DNA strand, and three left to right on the other.

The presence of multiple FAXC genes is a characteristic feature of invertebrate phyla that include Cnidaria and Mollusca. But the clustering of FAXC genes, with many or most of the multiple genes in the same genomic location, is not typical of the arrangement of the genes. Multiple FAXC genes are generally more dispersed throughout the genome, although there can be regions where two or three genes are in close proximity.

